# Suppressing microtubule detyrosination augments AAV2 endosomal escape and gene delivery

**DOI:** 10.1101/2025.08.12.669862

**Authors:** Shefali Tripathi, Shamshul Huda, Joydipta Kar, Dinesh Chandra, Jayandharan. R. Giridhara, Nitin Mohan

## Abstract

Adeno-associated virus (AAV) is a widely used vector for gene delivery, yet the host intracellular trafficking barriers often limit its efficacy. Here, we identify microtubule detyrosination—a tubulin post-translational modification—as a key regulator of AAV2 endo-lysosomal processing. Using super-resolution microscopy (SIM/STORM), we show that upon AAV2 endocytosis, the host upregulates microtubule detyrosination via GSK3β–CLASP2 signaling axis. Single particle tracking of the virus reveals that detyrosinated microtubules form a physical and functional barrier, restricting AAV2 motility and promoting lysosomal trapping. Restoring microtubule tyrosination—via tubulin-tyrosine ligase overexpression or pharmacological inhibition of detyrosination with parthenolide—boosted AAV2 endosomal escape, perinuclear accumulation, and gene delivery in cells. Notably, a clinically relevant prodrug of parthenolide, DMAPT, also displayed a similar trend of enhanced AAV2-driven factor IX expression in hemophilia B mouse models. Our findings uncover a host mechanism that reshapes the microtubule landscape to restrict AAV2 trafficking and identify microtubule detyrosination as a novel druggable target to improve AAV2-based gene therapies.

## Introduction

Viral vectors are at the forefront of gene therapy, enabling efficient delivery of genetic material into target cells ^1^. Among them, the adeno-associated virus (AAV) has emerged as a powerful tool valued for its diverse serotypes, high tissue tropism, and relatively low immunogenicity ^2,3^. While AAV holds therapeutic promise for genetic disorders like hemophilia B— caused by clotting Factor IX (FIX) deficiency, its efficacy is limited by critical bottlenecks. At low doses, protein expression is suboptimal; in contrast, higher doses risk activating immune responses ^4–6^. Current strategies, such as capsid engineering, aim to evade proteasomal degradation and minimize immune response. For instance, tyrosine-to-phenylalanine substitutions in the VP3 domain or alterations to SUMOylation/Neddylation sites enhance AAV2 capsid stability and transduction efficiency ^7–11^. However, intracellular trafficking barriers— particularly inefficient endosomal escape and lysosomal degradation— remain unresolved.

Studies on the AAV2 serotype have shown that AAVs enter cells via clathrin-mediated endocytosis, macropinocytosis, or CLIC/GEEC pathways ^12–15^. Post-cellular entry, AAV2 is sorted through the endo-lysosomal compartments. The acidic pH within late endosomes promotes AAV2 endosomal escape, where the capsid phospholipase A2 (PLA2) activity results in membrane rupture and the release of AAV2 into the cytosol ^16–18^. Subsequently, the nuclear localization signal (NLS) on the capsid facilitates AAV nuclear entry, allowing the capsid to uncoat, enabling gene expression ^19^. However, post-endosomal escape, most AAV2 particles are ubiquitinated and degraded in the cytoplasm, with only a fraction reaching the nucleus ^10,18,20^. Rapid retrograde transport on microtubules may mitigate this inefficiency by enabling endosomal escape near the nucleus, thereby evading cytoplasmic degradation. Supporting this notion, reports show that nocodazole-induced microtubule disruption impairs dynein-mediated AAV2 transport and gene delivery ^21–23^.

Microtubule post-translational modifications (PTMs)—such as acetylation, tyrosination, and detyrosination—form distinct microtubule subtypes within the cell, differentially regulating the activity of motor proteins and organelle trafficking ^24–26^. For example, detyrosinated microtubules influence lysosomal distribution and autophagosome-lysosome fusion ^27,28^. Intriguingly, viruses like HIV-1 and Influenza A modulate microtubule PTMs to facilitate entry or egress in immune and epithelial cells ^29,30^. Building on this evidence, we hypothesized that microtubule PTMs act as host-cell checkpoints governing AAV trafficking and infectivity.

In this study, we investigated the functional role and therapeutic potential of microtubule-PTMs in AAV-mediated gene delivery for hemophilia. We prioritized the AAV2 serotype for its hepatic tropism ^31^ and the established influence of microtubules on its cellular transport ^21^. Here, we show that AAV2 infection triggers microtubule detyrosination, which restricts viral motility and promotes its lysosomal entrapment. Conversely, suppressing detyrosination—via tubulin-tyrosine ligase overexpression or pharmacological inhibition with parthenolide—improves AAV2 retrograde trafficking and significantly enhances transduction efficiency in hepatic cells (Huh7). Notably, the clinically relevant parthenolide prodrug DMAPT recapitulates these effects in hemophilia B mice, boosting FIX expression. Our findings establish microtubule detyrosination as a druggable barrier to AAV2 gene delivery, offering a host-directed strategy to improve gene therapy outcomes.

## Results

### AAV2 endocytosis induces microtubule detyrosination via GSK3β-CLASP2 signaling

We investigated microtubule-PTM remodeling during AAV2 entry using a pulse-chase approach (see methods), where Huh7 cells exposed to AAV2 (2×10^4^ vgs/cell) were imaged with super-resolution structured illumination microscopy (SIM) at definite time points post-endocytosis (Figure. 1A-C). Endocytic AAV2 puncta increased progressively to 4 hours post-entry, followed by a decline by 8 hours—consistent with cytoplasmic degradation or nuclear translocation (Figure. 1A). Strikingly, detyrosinated microtubule levels mirrored this trend, peaking at 4 hours (mean=25 (3.8) %) and returning to baseline (mean= 10.5 (3.8) %) by 8 hours (Figure. 1B, D and Figure. S1A, S1B). Tyrosinated microtubule levels inversely correlated with detyrosination (Figure 1C, D), while acetylated microtubules remained undetectable (Figure S1C) with AAV2 endocytosis.

**Figure 1.**
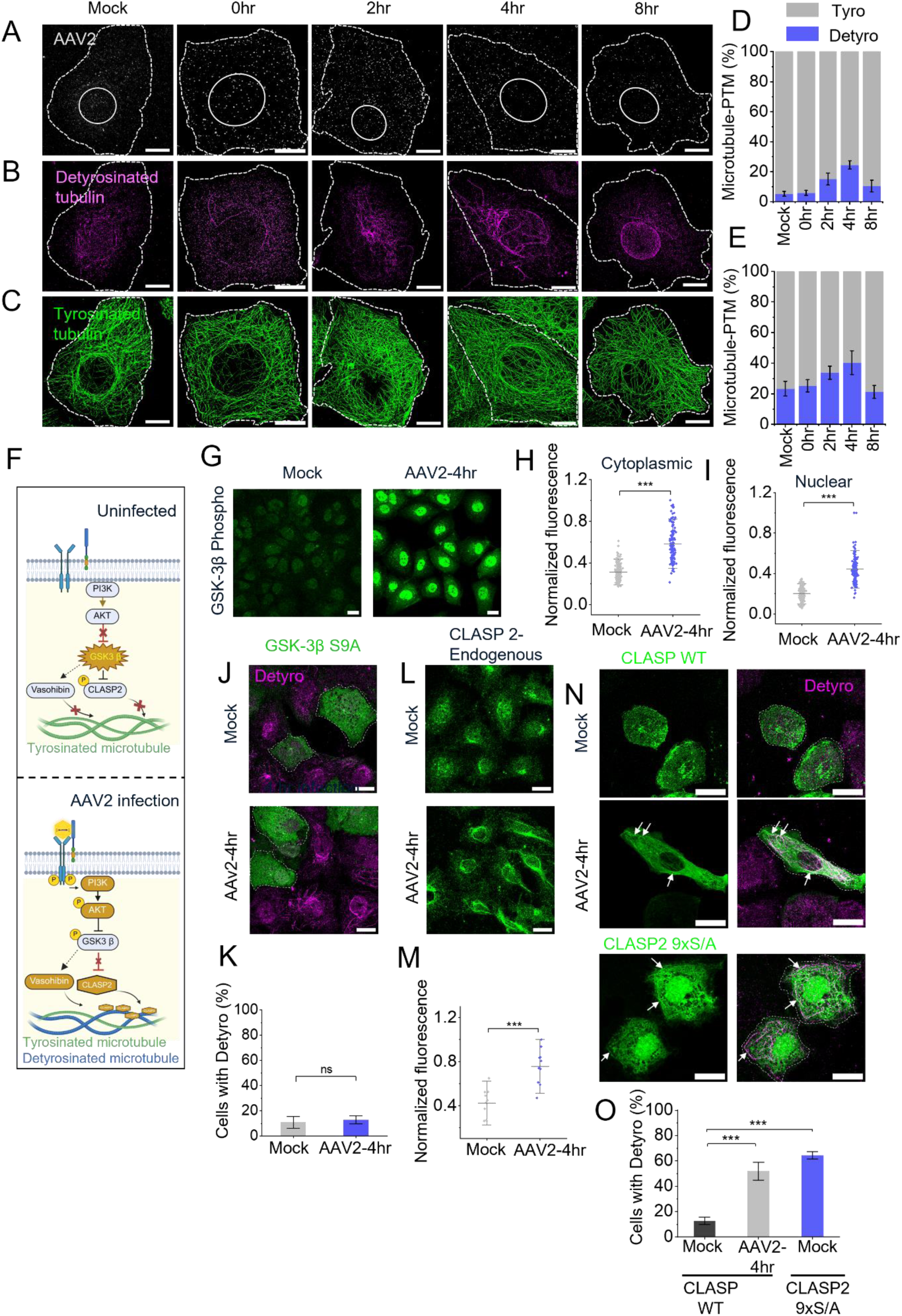
AAV2 endocytosis upregulates tubulin detyrosination via GSK3β-CLASP2 signaling. **(A–C)** Three-color SIM imaging of AAV2 (A), detyrosinated tubulin (B), and tyrosinated tubulin (C) in Huh7 cells at 0, 2, 4, and 8 hours post-AAV2 treatment. **(D–E)** Quantification of tyrosinated and detyrosinated microtubules in Huh7 (D) and BS-C-1 cells (E) across time points post-AAV2 treatment (mean ± SD; n = 15 cells analyzed across three independent experiments). **(F)** Schematic model of AAV2-induced signaling via RTK leading to GSK3β inhibition and CLASP2-mediated microtubule detyrosination. **(G–I)** Immunostaining (G) and quantification of cytoplasmic (H) and nuclear (I) phospho-GSK3β (Ser9) in mock– and AAV2-infected Huh7 cells at 4 hours, showing increased Ser9 phosphorylation (n=125 cells, analyzed across three independent experiments). **(J–K)** Expression of constitutively active GSK3β S9A mutant (green) and quantification of cells with detyrosinated microtubules (magenta) under mock or AAV2-infected conditions (n=190 cells, analyzed across three independent experiments). **(L–M)** Immunostaining (L) and quantification (M) of endogenous CLASP2, showing increased microtubule association in infected cells (in triplicate). Arrows indicate CLASP2 localized to microtubules. **(N–O)** Modulation of detyrosinated microtubules in cells overexpressing CLASP2 WT (green) or CLASP2 mutant-9xS/A (green) with AAV2 treatment compared to mock controls (n=160 cells, analyzed across three independent experiments). In the graphs the grey line or bar represents the mean, and whisker denotes the standard deviation. Significance was analyzed using two paired two-sample t-test, ***p<0.001 and “ns” indicating non-significant. Scale bar: (A-C) is 10 µm, (G, J, L and N) is 20 µm.

This dynamic was conserved in BS-C-1 cells, which exhibit higher baseline detyrosination (mean=23 (4.7) %). Here, AAV2 endocytosis induced a transient rise in detyrosination peaking at 4 hours (mean= 40.2 (7.7) %), followed by a decline to baseline (Figure 1E, S1D). Notably, detyrosination persisted in 75.7 (9) % of Huh7 cells at 4 hours but declined to 54.8 (11.2) % and 29.1 (3.6) % by 8 hours and 24 hours, respectively (Figure S1E and S1F).

We investigated how AAV2 entry increases microtubule detyrosination. Several co-receptors involved in AAV2 endocytosis are receptor tyrosine kinases (RTKs) ^32^, suggesting that AAV2 triggers RTK signaling cascade leading to detyrosination. RTK activation typically stimulates AKT phosphorylation downstream ^33^. Specifically, we investigated the RTK-AKT downstream kinase, glycogen synthase kinase 3 beta (GSK3β) ^34^, a known suppressor of the microtubule-stabilizing protein CLASP2 ^35,36^. Microtubule detyrosination mediated by carboxypeptidase, Vasohibin (VASH), is correlated with microtubule stability ^37,38^. We thus proposed that AAV2-triggered RTK activation leads to AKT-mediated phosphorylation of GSK3β, relieving its repression on CLASP2, thereby promoting CLASP2 microtubule binding and detyrosination by VASH (Figure 1F).

Supporting this model, AAV2 treatment markedly increased phospho-GSK3β levels compared to mock controls in Huh7 cells (Figure 1G-I). Further, a constitutively active, phosphorylation-resistant GSK3β mutant (S9A) blocked AAV2-induced detyrosination, indicating that rather than a direct mechanism, AKT promotes detyrosination indirectly by inhibiting GSK3β (Figure 1J and 1K). Notably, the same results were seen in BSC-1 cells, suggesting this pathway is conserved (Figure S1G). We next assessed CLASP2, which showed an increased microtubule association following AAV2 endocytosis compared to mock controls. (Figure 1L and 1M). Further, while the wild-type CLASP2 overexpression induced detyrosination only in the AAV2 treated cells and colocalized with detyrosinated tracks (Figure. 1N-O and Figure. S1H), a GSK3β-insensitive CLASP2 mutant (9xS/A) was sufficient to promote detyrosination even without AAV2 (Figure. 1N-CLASP2 9xS/A and 1O), highlighting GSK3β as a key negative regulator of CLASP2-mediated detyrosination. Overall, our results reveal that AAV2 activates RTK–AKT signaling, leading to GSK3β inhibition, promoting CLASP2 microtubule stabilization and detyrosination by VASH. However, GSK3β may also regulate VASH-induced detyrosination through CLASP2-independent mechanisms, which warrants further investigation.

### AAV2 accumulates on detyrosinated microtubules and exhibits reduced motility

As previous results indicated that AAV2 infection promotes microtubule detyrosination via host signaling pathways, we next asked whether this post-translational modification affects the spatial distribution and motility of AAV2. Furthermore, with the help of SIM imaging and quantification (Figure S2A), we observed a time-dependent accumulation of AAV2 particles on detyrosinated microtubules in both Huh7 (Figure 2A-C) and BS-C-1 cells (Figure S2B-D). Super-resolution STORM imaging further confirmed the localization of AAV2 on detyrosinated microtubules with ∼20 nm precision (Figure S2E and S2F). These findings demonstrate that AAV2 endocytosis induces a ∼25% increase in detyrosinated microtubules, leading to the accumulation of ∼35% of AAV2 and a ∼1.4-fold enrichment on detyrosinated microtubules at 4 hours (Figure 2C), suggesting that detyrosination acts as a roadblock to AAV2 retrograde transport. Furthermore, the subsequent decline in cytoplasmic AAV2 at 8 hours indicates that detyrosination likely promotes AAV2 degradation.

**Figure 2.**
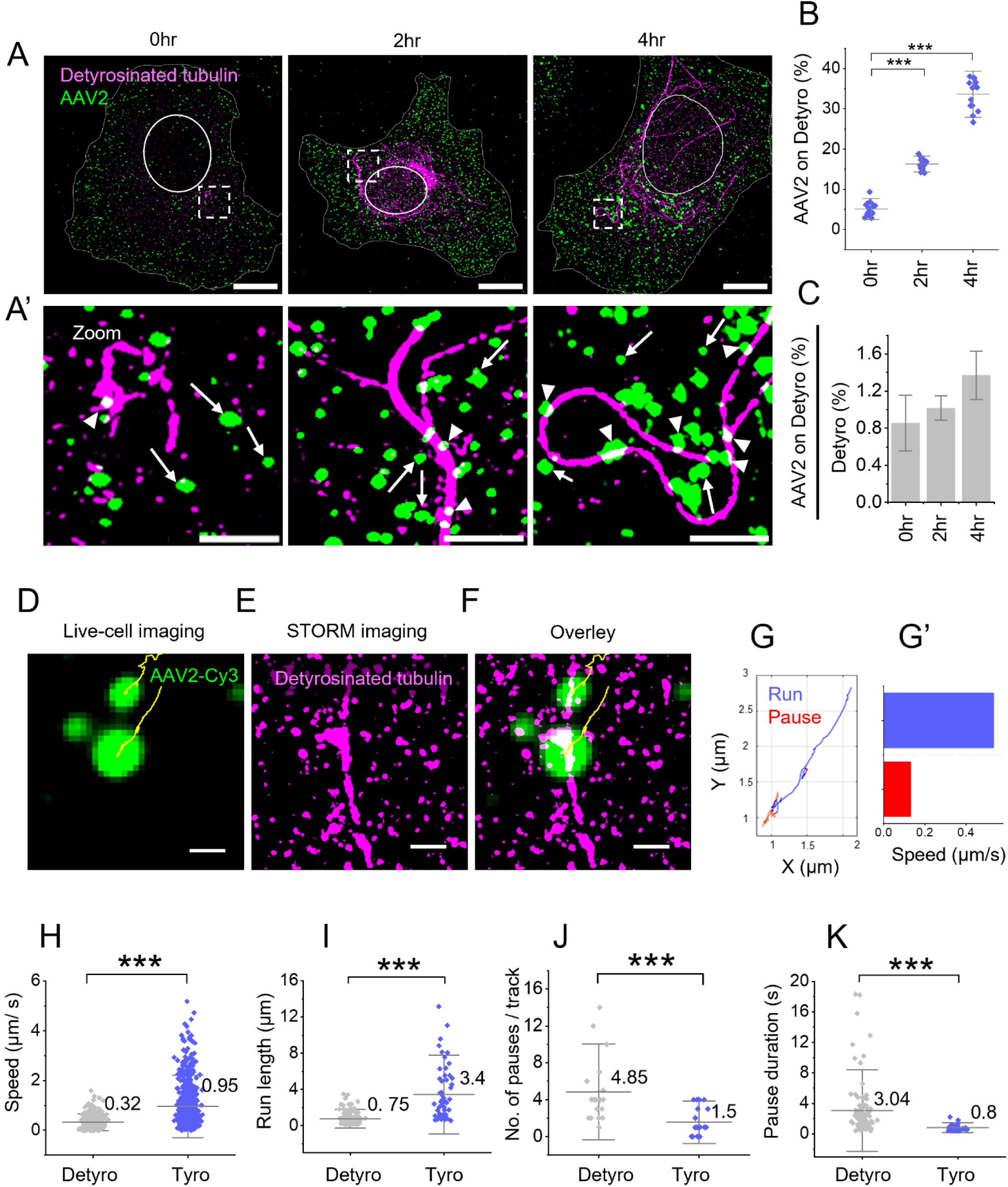
Detyrosinated Microtubules Influence AAV2 Distribution and Motility Dynamics. **(A-A’)** Two-color SIM images of Huh7 cells treated with AAV2 (green) and immune stained for detyrosinated microtubules (magenta) at 0-, 2-, 4-hour. Zoom of the respective insets (A’) given in (A). Arrowhead and arrows show AAV2 localized to and not localized to detyrosinated microtubules, respectively. **(B-C)** Quantification of the localization (B) and enrichment (C) of AAV2 on detyrosinated microtubules at different time points post-infection in Huh7 cells (10 cells analyzed for each time point; n=10). **(D-F)** Correlative live-cell and STORM imaging of AAV2 and detyrosinated microtubules in BS-C-1 cells. Trajectory of AAV2 motility (yellow) from the live-cell imaging (D). STORM image of the detyrosinated microtubules (magenta) from the same region (E). AAV2 trajectory from A overlaid with STORM image of detyrosinated microtubule from E (F). **(G-G’)** Color-coded AAV2 trajectory shows processive run phases in blue and non-processive pause phases in red (G) and their corresponding speed measurements (G’). **(H-K)** Quantitative analysis of speed (H), run length (I), number of pauses per track (J), and fraction of time spent pausing (pause duration) (K) of AAV2 motility on detyrosinated tubulin compared to tyrosinated tubulin (20 tracks analyzed per condition: n=20). In the graphs the grey line represents the mean and whisker denotes the standard deviation. Significance was analyzed using two paired two-sample t-test, ***p<0.001. Scale bar: (A) is 10µm, (A’) is 2 µm and (D-F) is 1 µm.

To resolve whether detyrosination impedes AAV2 retrograde transport, we used the correlative live-cell and STORM imaging approach ^27,28,39^. At 1-hour post-endocytosis, we tracked Cy3-tagged AAV2 in live cells for ∼ 1 minute, followed by *in situ* immunostaining and STORM imaging of detyrosinated microtubules within the same cell. By mapping AAV2 single-particle trajectories (SPT) onto the STORM image (Figure 2D-E) and segmenting processive (run) and non-processive (pause) states (Figure 2G and 2G’), we quantified AAV2 motility, using a custom SPT analysis algorithm ^39^. Our results reveal that AAV2 transitions from run to pause states (Figure 2F-G’, Figure S2G; supporting videos 1 & 2) or remains static (confined motion) on detyrosinated microtubules (Figure S2H; supporting videos 3). In contrast, it moved unimpeded when not moving on detyrosinated microtubules (Figure S2I, supporting video 4).

Quantitative analysis demonstrated that detyrosinated microtubules reduced AAV2 speed by 66% (mean= 0.32 (0.08) µm/s vs. 0.95 (0.12) µm/s on tyrosinated tracks, Figure. 2H) and shortened run lengths by ∼78% (0.75 (0.21) µm vs. 3.4 (0.45) µm; Figure. 2I). Detyrosination also increased pause frequency (mean= 4.85 (0.9) pauses/track vs. 1.5 (0.3), Figure. 2J) and prolonged pause durations (mean= 3.04 (0.6) s vs. 0.8 (0.2) s, Figure. 2K). Together, these data demonstrate that detyrosination acts as a physical barrier to AAV2 motility, either by trapping it or intermittently obstructing its movement, potentially limiting its retrograde transport and nuclear entry.

### Suppressing tubulin detyrosination accelerates AAV2 transport towards the nucleus

Next, we modulated detyrosination/tyrosination levels to assess their impact on AAV2 motility and cytoplasmic distribution. Overexpression of the tubulin-detyrosination enzyme vasohibin (VASH) increased detyrosinated microtubules in Huh7 cells (Figure S3A) and reduced AAV2 motility, whereas overexpression of the tyrosination enzyme tubulin tyrosine ligase (TTL) increased the tyrosination/ detyrosination state of the microtubules (Figure S3A) enhanced AAV2 displacement compared to untreated controls (Figure 3A and 3B, supporting video 5, 6 and 7). Additionally, parthenolide, a well-known VASH inhibitor, completely suppressed microtubule detyrosination (at 20 µM, see Methods and Figure S3B and S3C) and significantly increased AAV2 motility (Figure 3A and B, Supporting video 8). Additionally, AAV2 exhibited higher speed and run length in parthenolide-treated cells in comparison to the untreated control cells (Figure S3D-F).

**Figure 3.**
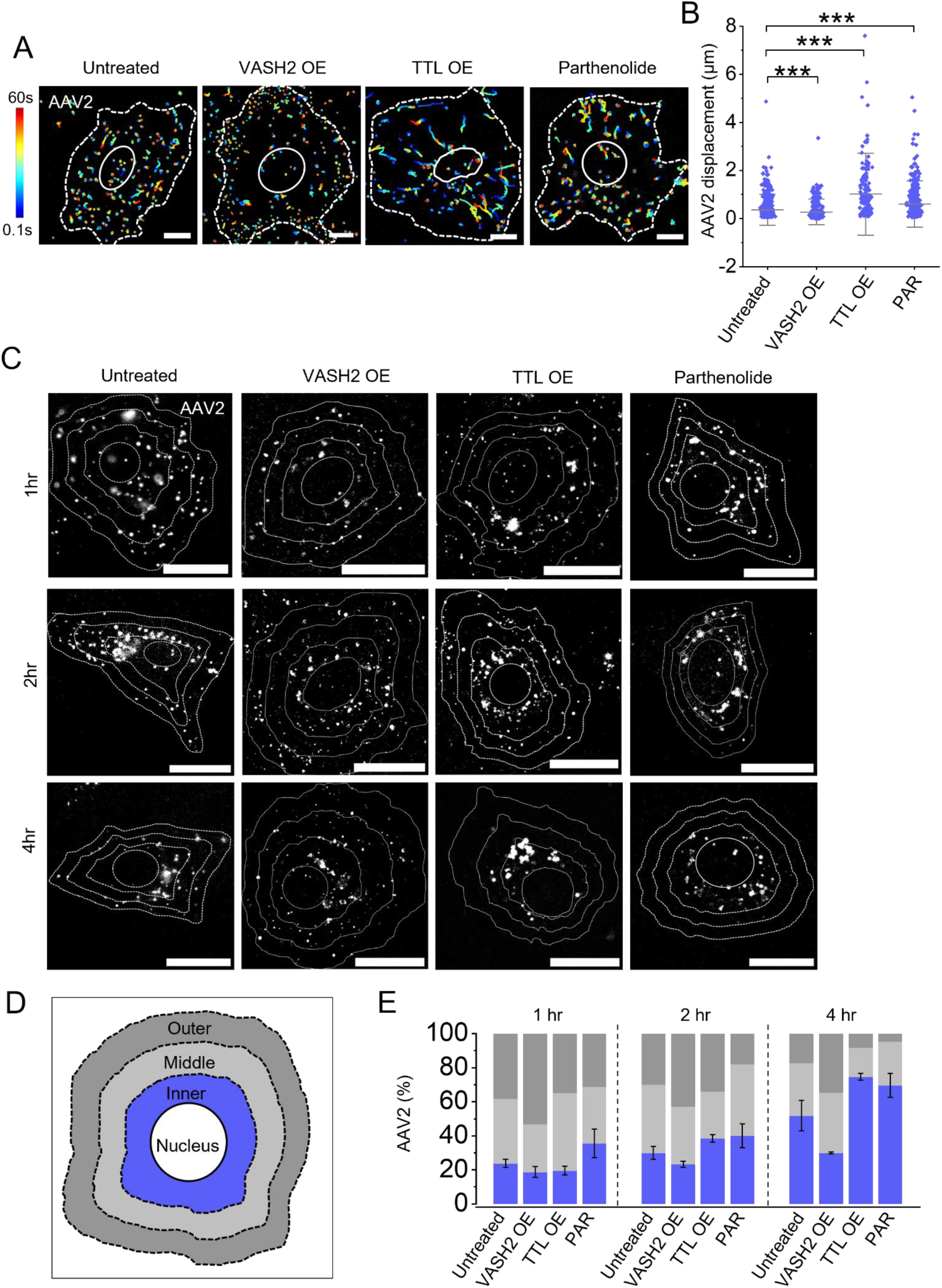
Suppression of detyrosinated microtubules improves AAV2 motility and perinuclear accumulation. **(A-B)** Live cell imaging and track displacement measurement of Cy3 tagged AAV2 in Huh7 cells in untreated control conditions and cells overexpressing VASH2-GFP or TTL-GFP or pretreated with parthenolide. Color-coded trajectories indicate blue for the start and red for the end of the tracks acquired for 60 seconds duration. (B) Quantification of AAV2 track displacement obtained from movies in A. Whiskers represent the standard deviation, and the grey line shows the mean for ∼500 trajectories from n=4 cells analyzed for each condition. **(C-E)** Time dependent spatial distribution analysis of AAV2 in Huh7cells. TIRF imaging of AAV2-Cy3 (white) in cells fixed at 1-, 2-, and 4-hour after AAV2 endocytosis (C). Schematic representation of the division of the cytoplasmic region into three equal zones: outer (near plasma membrane), inner (perinuclear) and middle (between zone 1 and 2), to analyze the spatial distribution of AAV2 within cells (D). Percentage of AAV2 distribution across the zones over time in Huh7 cells (E). Bars represent the mean and whisker represent the standard deviation for n= 10 cells analyzed in each condition. Statistical significance assessed using a two paired two-sample t-test, with ***p<0.001. Scale bar: (A) 10 µm and (C) 20 µm.

To examine the spatial distribution of AAV2 under variable tyrosination/detyrosination levels, we divided cells into three zones: inner (perinuclear), middle (mid-cytoplasmic), and outer (near the plasma membrane) (see Methods, Figure 3D). We then quantified AAV2 localization in these zones over time. VASH2 overexpression restricted most AAV2 to the outer zone, with only a fraction (mean= 28.25 (5) % at 4 hours) reaching the perinuclear zone. In contrast, TTL overexpression facilitated the accumulation of a larger pool (mean= 75 (0.6) % at 4 hours) of AAV2 near the nucleus (Figure 3C and E). Similarly, parthenolide-mediated suppression of detyrosination significantly increased AAV2 accumulation in the inner zone (mean= 69.6 (7) % at 4 hours). These findings indicate that detyrosinated microtubules impede AAV2 retrograde transport to the nucleus, while suppressing detyrosination or enhancing tyrosination accelerates perinuclear accumulation, potentially improving AAV2 nuclear entry.

### Microtubule tyrosination enhances AAV2 endosomal escape

To determine how the tyrosination/detyrosination status of microtubules regulates AAV2 endosomal sorting, we tracked over time the colocalization of AAV2 with Rab5 (early endosomes), Rab7 (late endosomes), and LAMP1 (lysosomes) in parthenolide-treated versus untreated cells. Parthenolide treatment significantly reduced AAV2 retention in early endosomes (Figure S4A and S4B), indicating a rapid sorting of AAV2 from early to late endosomes. Notably, unlike the progressive accumulation of AAV2 in late endosomes and lysosomes observed in control cells, parthenolide treatment accelerated AAV2 exit from late endosomes (Figure 4A and 4B) and reversed its accumulation in lysosomes (Figure 4C and 4D), suggesting microtubule tyrosination enhanced AAV2 endosomal escape and diverted it from lysosomes.

**Figure 4.**
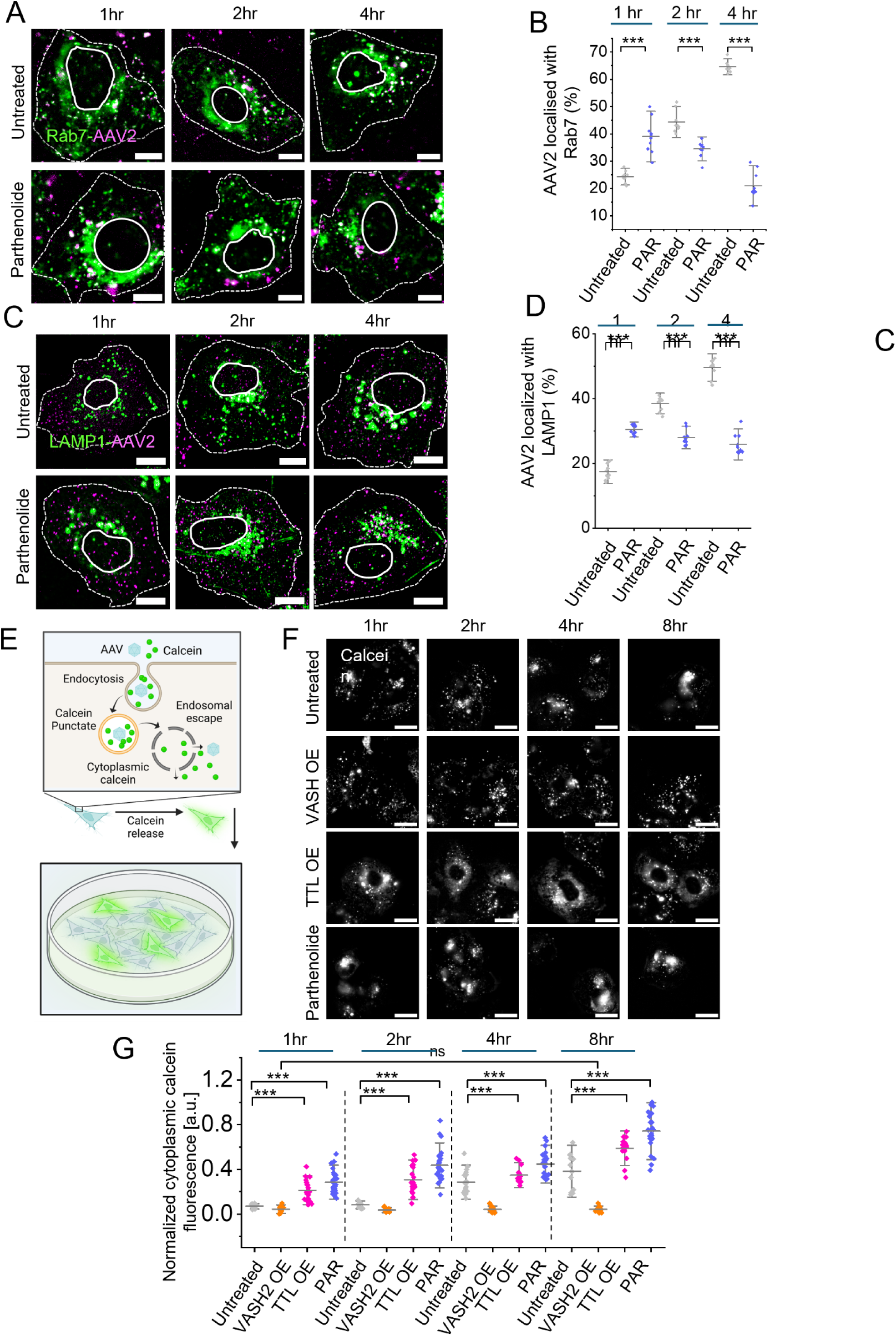
Tyrosinated microtubules promotes AAV2 endosomal escape. **(A-B)** Time dependent colocalization analysis of AAV2 with late endosomes in Huh7 cells. Confocal imaging of Rab7-GFP (green) along with Cy-3 tagged AAV2 (magenta) at different time points in Huh7 cells in untread conditions and pre-treated with parthenolide (A). Percentage of AAV2 colocalized with Rab7 decorated late endosomes over time (n=10) (B). **(C-D)** Confocal imaging and Quantification of AAV2 (magenta) colocalization with lysosomes decorated by LAMP1 (green) (C). While untreated control cells show significant accumulation of AAV2 in lysosomes over time, parthenolide-treated cells show no significant change in lysosomal colocalization with time (n=10) (D). **(E-G)** AAV2 endosomal escape measurement using calcien fluorescence assay in Huh7 cells. Schematic representation of calcein fluorescence assay. Endosomal calcein fluorescence appears as a punctate signal whereas, AAV2 mediated endosomal rupture leads to diffused cytoplasmic calcein fluorescence (E). Live-cell imaging of calcein (white) in AAV2 infected cells at 1-, 2-, 4-, and 8-hour post AAV2 endocytosis in untreated control, VASH2-GFP overexpressed, TTL-GFP overexpressed, or parthenolide pretreated conditions (F). Quantification of cytoplasmic calcein fluorescence over time (n=22) (G). In the graphs, whiskers show the standard deviation, the grey line indicates the mean value. Statistical significance was determined using a two paired two-sample t-test, with ***p<0.001 indicating significance and “ns” indicating non-significance. Scale bars: (A-C) 10 µm and (D) 20 µm.

To directly assess endosomal escape facilitated by tyrosinated microtubules, we employed the calcein green assay (see Methods). Calcein, a membrane-impermeable dye, enters cells via endocytosis and its punctate fluorescence (confined to intact endosomes) transitions to a diffuse cytoplasmic signal upon AAV2-induced membrane rupture (Figure 4E). In cells overexpressing VASH2 (promoting detyrosination), calcein remained punctate until 8 hours post-entry, indicating minimal escape (Figure 4F and 4G). Conversely, TTL overexpression (inducing tyrosination) or parthenolide treatment triggered calcein cytoplasmic diffusion within 1 hour, with signal intensity increasing over time, demonstrating robust AAV2 escape (Figure 4F and 4G). These findings suggest that suppressing detyrosinated microtubules through TTL overexpression or parthenolide treatment prevents lysosomal entrapment and promotes AAV2 endosomal escape, likely facilitating its nuclear translocation.

### Suppression of microtubule detyrosination enhances AAV2 transduction efficiency

While endosomal escape is essential for AAV2 nuclear entry, it exposes the virus to ubiquitination and proteasomal degradation. To determine whether suppressing microtubule detyrosination preferentially promotes AAV2 nuclear entry over degradation, we quantified transduction using an AAV2 vector with a GFP reporter transgene. Overexpression of VASH2 (inducing detyrosination) reduced GFP-positive cells to 14.6 (2.8) %, whereas TTL overexpression (promoting tyrosination) markedly increased transduction to 82.3 (1.5) % (Figure 5A and 5B). AAV2 EGFP expression inversely correlated with VASH2 levels but increased dose-dependently with TTL (Figure S5A and S5B), confirming that detyrosination impedes AAV2 nuclear translocation.

**Figure 5.**
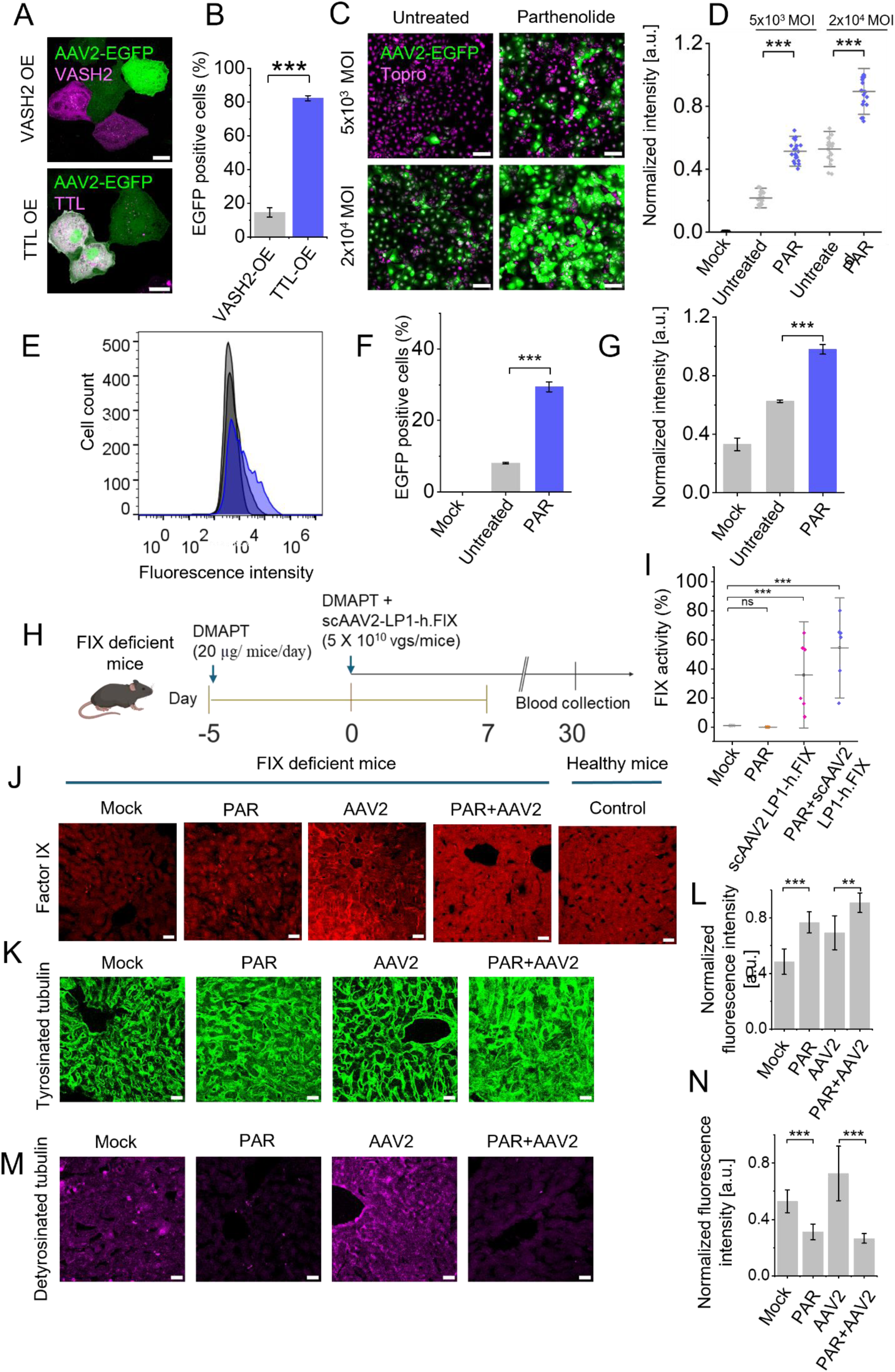
Detyrosinated microtubule modulation and combinatorial parthenolide+AAV2-FIX therapy in cellular and animal models. **(A-B)** Assessment of the impact of VASH and TTL overexpression on AAV2 transduction efficiency. Confocal imaging of Huh7 cells overexpressing VASH2 (A-upper panel, magenta) or TTL (A-lower panel, magenta) infected with AAV2 EGFP, where expression of EGFP transgene (green) indicates AAV2 transduction (A). Percentage of EGFP-positive cells among VASH2-or TTL-overexpressing populations, highlighting their influence on transduction. n=100 cells analyzed for each condition from triplicate (B). **(C-D)** Huh7 cells, untreated or pretreated with parthenolide, were infected with AAV2-EGFP at low (MOI:5×10³ vg/cell) and high (MOI: 2×10⁴ vg/cell) vector doses. Confocal imaging shows GFP expression (green) in infected cells, with topro staining nuclei (magenta) (C). Fluorescence intensity quantification confirmed enhanced AAV2-EGFP expression in parthenolide-treated cells at both MOIs (no. of images; n=18) (D). **(E-G)** Flow cytometry analysis (E) and quantification of GFP-positive cells (F) and normalized fluorescence intensity (MOI: 5×10³ vg/cell) (G), also indicate improved transduction efficiency with parthenolide treatment n=10000 cells were analyzed by flow cytometry for each condition in triplicate. **(H-J)** Schematic illustrates the *in-vivo* therapeutic evaluation of parthenolide combined with scAAV2-LP1-h.FIX in a hemophilia B mouse model (H). Parthenolide was administered from day –5 to day –1, followed by AAV2-FIX administration on day 1 and blood collection on day 30 for FIX activity analysis (no. of mice/group; n=5 to 7). Chromogenic assay (I) compared FIX activity across four groups: mock (untreated hemophilia B mice), PAR (parthenolide only), AAV2 (vector without drug), and PAR+AAV2 (combination therapy). Immunohistochemistry and confocal imaging of liver sections showed FIX expressions (red) in the PAR+AAV2 group compared to controls and healthy C57 mice (J). **(K-N)** Quantification of tyrosinated (K-L) and detyrosinated tubulin (M-N) in liver sections further revealed differences across groups, highlighting the impact of combination therapy. In graphs, the grey line represents the mean value, and whiskers indicate the standard deviation. Statistical significance was determined using two paired two-sample t-tests, with ***p<0.001 indicating significance. For *in-vivo* FIX expression significance was analyzed using two-way ANOVA, Scale bar: (A, J, K and M) 20 µm and (C) 100 µm.

**Figure 6.**
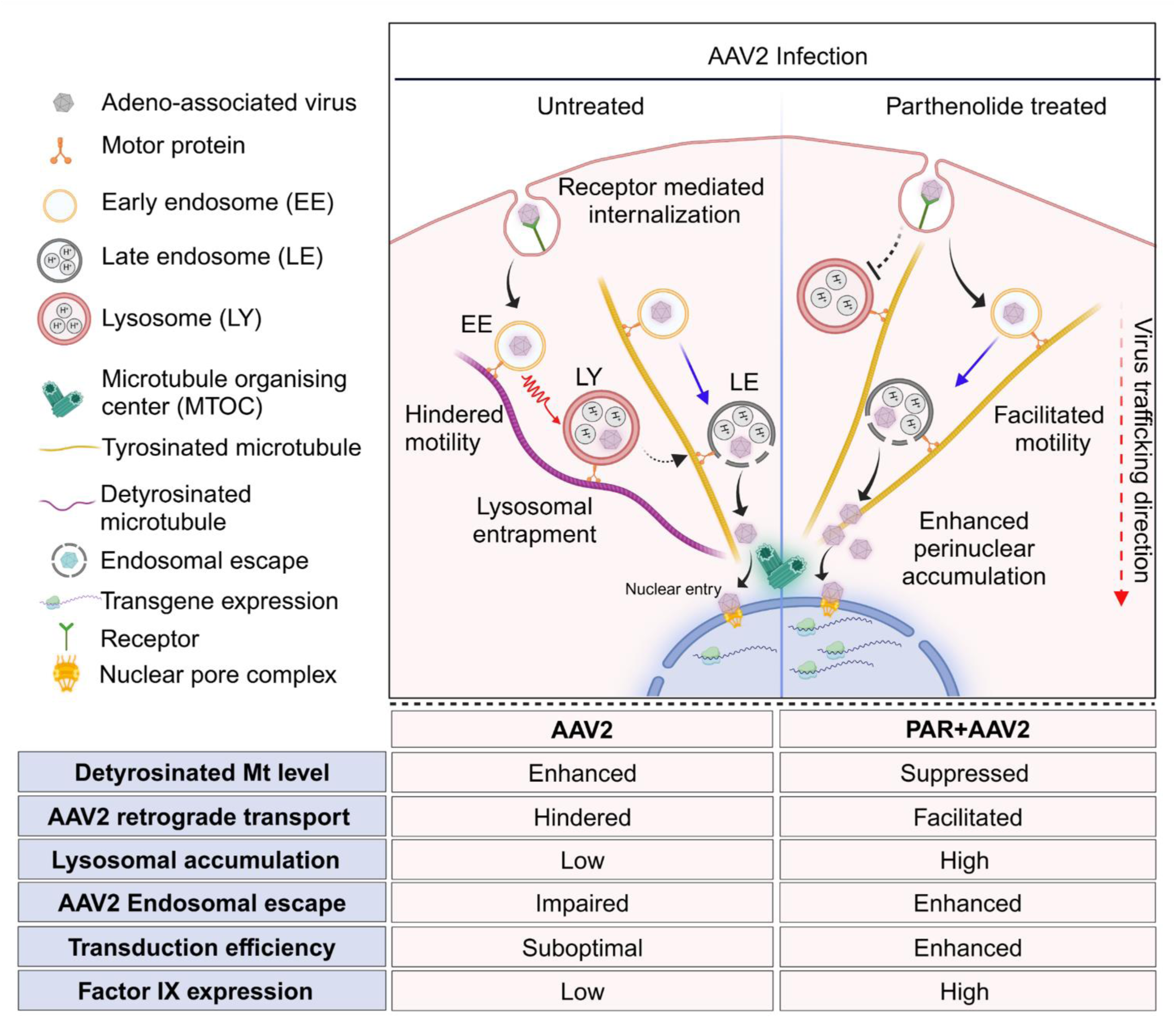
Proposed model. The Figureure illustrates AAV2 intracellular transport emphasizing the role of microtubule post-translational modifications in regulating AAV2 trafficking. It highlights the hindered motility of AAV2 on detyrosinated microtubules (red zigzag arrows), leading to its accumulation in lysosomes, with minimal endosomal escape. Conversely, parthenolide treatment (right panel) suppresses detyrosinated tubulin, enabling accelerated transport (blue arrows), increased endosomal escape close to the nucleus, leading to enhanced transduction efficiency.

Pharmacological suppression of detyrosination with parthenolide mirrored these effects. Parthenolide-pretreated cells exhibited a ∼2-fold increase in GFP fluorescence intensity and a ∼1.5-fold higher percentage of GFP-positive Huh7 cells compared to untreated controls, across viral doses (5 × 10³ or 2 × 10⁴ vgs/cell; Figure 5C and 5D). Consistently, flow cytometry analysis revealed 3.6-fold more GFP-positive Huh7 cells (mean= 29.4 (0.91) % vs. 8 (0.16) % in controls; p ≤ 2.4 × 10⁻⁶) and ∼1.5-fold higher GFP intensity following parthenolide treatment (Figure 5E-G), demonstrating enhanced AAV2 transduction with pharmacological detyrosination suppression *in vitro*.

To validate the therapeutic relevance of suppressing detyrosination as a strategy to improve AAV2-mediated gene therapy *in vivo*, we used hemophilia B mouse model (Figure 5H, see Methods). We used pro-drug Dimethylamino Parthenolide (DMAPT) due to its bioavailability and prior use in suppressing detyrosinated microtubule in animal models ^40^. Mice receiving scAAV2-LP1-h.FIX vector (5 × 10¹⁰ vgs/mouse) pretreated with DMAPT showed an elevated FIX coagulant activity (mean= 54.47 (22.9) %, Figure 5I) compared to mice without DMAPT pretreatment (mean= 35.84 (24.33) %, Figure 5I). Additionally, in liver tissues of DMAPT-treated mice, FIX expression increased. (Figure 5J). Immunohistochemistry reveals the reduced microtubule detyrosination over tyrosination (Figure 5 K-N) further supporting the role of detyrosination suppression in enhancing AAV2 efficacy.

Thus, suppressing microtubule detyrosination—a host-cell barrier that traps AAV2 on retrograde transport pathways—enhances viral nuclear entry and gene delivery efficacy, offering a clinically actionable strategy to optimize AAV-based therapies.

## Discussion

Our study uncovers a new role for detyrosinated microtubules in cellular defense against AAV2 infection in liver cells and tissues. Upon the AAV2 infection, the host cell increases the detyrosinated microtubule levels to impede AAV2 trafficking towards the nucleus, limiting its transduction. Suppressing microtubule detyrosination enhances AAV2-mediated factor IX gene transduction efficiency in liver cells and hemophilia B mouse models, offering a promising strategy for augmenting AAV-mediated gene therapy.

Viruses often hijack host cytoskeletal pathways to facilitate their intracellular transport. For example, HIV-1 promotes the formation of stable microtubules ^29^ via the Diaphanous-related formin (DRF) proteins Dia1 and Dia2 ^44^, while microtubule-associated proteins (MAPs) such as CLASP2 support its intracellular motility ^42^. Intriguingly, Wnt/β-catenin-GSK3β signaling is reported to modulate HIV replication ^45^. Similarly, Influenza A virus infection increases microtubule acetylation ^30^, and Tubacin mediated suppression of acetylation reduces viral replication ^46^. Notably, RTK/AKT pathways are activated during Influenza A infection as well, promoting anti-apoptotic responses ^47^, and likely facilitating cytoskeletal remodelling for viral trafficking.

Our data reveal that AAV2 infection engages a related but distinct pathway. Specifically, AAV2 internalization activates RTK–AKT–GSK3β signaling, which in turn elevates detyrosinated microtubule levels. Notably, we show that this detyrosination is not a direct effect of viral components but rather a downstream consequence of host signaling responses triggered by AAV2 entry. Unlike HIV-1 or Influenza A, where stable acetylated or detyrosinated microtubules promote infection, AAV2 is impaired by detyrosinated microtubules, suggesting that non-enveloped viruses face distinct cytoskeletal barriers compared to enveloped viruses.

Previous studies show that nocodazole-induced microtubule depolymerization impairs AAV2 endosomal escape and infectivity ^21^. Nocodazole selectively targets labile microtubules at micromolar concentrations ^41,43^, suggesting AAV2 retrograde transport is dependent on labile tyrosinated microtubules. Consistently, our findings show that increasing tyrosinated microtubule pools enhances AAV2 perinuclear trafficking and transduction, in contrast to HIV-1 and Influenza A, which exploit stable microtubule populations.^29,30,44,46^. This divergence may stem from differences in viral entry receptors and the distinct mechanisms employed by enveloped and non-enveloped viruses to exploit the cytoskeleton to their advantage for endosomal sorting and trafficking to evade cellular degradation.

Mechanistically AAV2 enters cells via Heparan Sulfate Proteoglycan (HSPG), mediated endocytosis with growth factor receptors acting as co-receptors ^48–51^. Following internalization, HSPG is directed to lysosomal degradation, a process disrupted by vinblastine-induced microtubule depolymerization ^52^. Notably, vinblastine has been shown to depolymerize stable microtubules marked by acetylation and impair autophagosome-lysosome fusion ^24^. Vinblastine is likely to affect all stable microtubules, including detyrosinated microtubules, suggesting that upon AAV2 internalization, HSPG is directed to lysosomes along the detyrosinated microtubules. Furthermore, previous reports show that lysosomal motility is hindered on detyrosinated microtubules, enriching lysosomes on detyrosinated microtubules, facilitating autophagosome-lysosome fusion ^28^. Taken together, these observations suggest that detyrosinated microtubules foster a lysosomal degradation hub where cellular cargoes and viral particles from endocytic and autophagic pathways accumulate proximal to lysosomes, facilitating their fusion and degradation. Congruently, we observe detyrosinated microtubules accumulate AAV2, and their suppression disrupts AAV2 lysosomal entrapment.

Parthenolide and its prodrug, DMAPT, are promising inhibitors of microtubule detyrosination ^40,53,54^. While parthenolide’s anticancer effects stem from NF-κB modulation ^55^, its detyrosination inhibition at micromolar concentrations ^38^ is independent of NF-κB activity ^53^. Parthenolide promotes axon regeneration in injury-induced mouse models ^40,54^ and reduces muscle stiffness by modulating detyrosinated microtubule, offering therapeutic potential for muscular dystrophies ^56^ and heart failure ^57^. In our hemophilia B mouse model, DMAPT enhanced AAV2-mediated human FIX gene delivery. These findings expand DMAPT’s potential applications beyond cardiac disease and cancer, highlighting its role in optimizing gene therapy outcomes.

Conclusively, we propose that microtubule-PTMs regulate AAV2 trafficking and transgene expression (Figure 7). AAV2 enters cells via receptor-mediated endocytosis ^13–15^, and traffics through early and late endosomes ^17,58^ along microtubules ^21^. Endosomal acidification activates the capsid phospholipase A2 domain, enabling membrane rupture and viral escape ^16^, leading to nuclear entry for gene expression ^59^. We demonstrate that AAV2 transport towards the nucleus is impaired on detyrosinated microtubules, causing lysosomal accumulation and limiting endosomal escape, resulting in reduced transgene expression. In contrast, parthenolide treatment promotes rapid AAV2 transport along tyrosinated microtubules, enhancing endosomal escape near the nucleus, preventing cytoplasmic degradation, and boosting nuclear entry and transgene expression. Finally, these findings highlight the potential of a drug-vector co-administration strategy (co-administering parthenolide with AAV2) to enhance transgene expression and improve therapeutic outcomes for hemophilia B patients.

Our study is limited to the AAV2 serotype, and the effects of detyrosination suppression should be validated across other AAV serotypes. The immune response to pharmacological modulation requires further investigation. Nonetheless, our findings establish microtubule detyrosination as a key regulator of AAV2 trafficking and present a novel target for optimizing AAV-mediated gene therapy.

## Material and methods

### AAV2 production and dye conjugation

AAV2 was prepared and produced as described earlier ^7,8,60^. Briefly, recombinant AAV2 (scAAV2) vectors containing human factor FIX (h.FIX) gene driven by LP1 promoter/enhancer were generated by triple transfection protocol as described previously ^60^. Purified vectors were quantified by quantitative PCR using polyA region specific forward and reverse primers after DNase treatment ^61^. For live imaging of AAV2, dye and virus were conjugated as described previously ^12^ with some modifications. Briefly AAV2 (3×10¹² vgs; 20 µg) was incubated with Cy3 dye (Cytiva PA23001) (50 µM) in a conjugation buffer (0.1 M sodium bicarbonate, pH 9.3; Sigma Aldrich) for 8 hours at 4°C. The buffer was prepared by dissolving NaHCO₃ in water and adjusting the pH to 9.3 by gradually adding 0.1M Na₂CO₃ while stirring. To remove unbound dye, the mixture was subjected to centrifugal filtration using 0.5 ml filters (Merck Millipore) in a dialysis buffer (10% glycerol, 10 mM Tris, and 50 mM NaCl, pH 7.4 in deionized water). Centrifugation was performed at 14,000 g for 15 minutes at 4°C, and the filtration process was repeated approximately 8 times or until the solution appeared visibly clear, confirming the successful removal of unbound dye. The virus was then aliquoted and stored in –80°C. The degree of labelling of Cy3-AAV2 was determined by spectrophotometry and it was speculated that the dye reacted with the ε-amine of lysine residues and the hydroxyl groups of serine and threonine. The labelled virus titer was determined by quantitative PCR.

### Cell culture and transfection

Human hepatoma-derived Huh7 (kind gift from S. Das; Indian Institute of Science, Bengaluru, India) were maintained in Iscove’s Modified Dulbecco’s Medium (IMDM, Gibco) and African green monkey kidney epithelial cells, BSC1 (kind gift from Melike Lakadamyali, University of Pennsylvania, USA) were maintained in Minimum Essential Medium (MEM, Gibco) supplemented with 2mM L-Glutamine, 1mM Sodium pyruvate. Media were supplemented with 10% fetal bovine serum (FBS, Hyclone, USA), and penicillin-streptomycin (Invitrogen) or Anti-Anti (Gibco). All the cells were maintained at 37 ^0^C and 5% CO_2_. For imaging experiments, cells were seeded in 4 well-chambered, 8 well-chambered, or single-well imaging dishes (Ibidi, Gräfelfing, Germany) at ∼10^4^ cells per well with 200-400μl media. Plasmid transfection was performed using X-Treme GENE HP DNA transfection reagent (Sigma Aldrich). Cells were transfected with 300 ng DNA for 24 hours. pShuttle-mRFP-GSK3 S9A was a gift from Torsten Wittmann (Addgene plasmid # 24371), pAd/CMV/V5-Clasp2α was a gift from Torsten Wittmann (Addgene plasmid # 89763), pEGFP-CLASP2 512-650 9xS/A was a gift from Torsten Wittmann (Addgene plasmid # 24410), Rab7-GFP, Rab5-GFP, and LAMP1-GFP plasmids were a kind gift from Indranil Banerjee (IISER Mohali, India), VASH2-Mcherry and TTL-Scarlet, were a kind gift from Carsten Janke (Institut Curie, France) and VASH2-GFP, and TTL-GFP were kindly provided by Minhaj Sirajuddin (InStem, Bengaluru, India).

### Pulse-chase treatment

For pulse-chase experiments, cells were incubated with adeno-associated virus serotype 2 (AAV2) at a multiplicity of infection (MOI) of 2×10^4^ (2×10^4^ vector genomes ‘vgs’ / cell) in serum-free media at 37°C for 2 hours to facilitate viral adsorption onto the cell surface ^62,63^. Following incubation, unbound AAV2 was removed by washing the cells with phosphate-buffered saline (PBS). Cells were then incubated in media supplemented with 10% fetal bovine serum (FBS) at 37°C for chase time points of 0, 2, 4, and 8 hours. Mock-treated control cells underwent the same procedure without AAV2 exposure.

### Parthenolide treatment

Cells were incubated with varying concentrations of parthenolide for 48 hours, and cell viability was assessed using the MTT assay (see Figureure S4B) to determine the half-maximal inhibitory concentration (IC_50_) of parthenolide (P0667, Sigma Aldrich) to be 100 µM. Additionally, Huh7 cells were treated with different concentrations of parthenolide to identify the optimal dose for suppressing detyrosinated microtubule, which was found to be 20 µM (see Figureure S4C). Further in all the experiment cells were pretreated with 20 µM parthenolide for 2hr before the AAV2 treatment.

### Endosomal escape with calcein

Cells were either untreated or pretreated with 20 µM parthenolide for 2 hours. Subsequently, cells were treated with AAV2 for 2 hours in serum free medium as mentioned in pulse chase treatment above. Immediately after media change, calcein was added at a final concentration of 100 µg/ml in FBS containing medium, marking the 0 hour time point. Further cells were imaged at 0-hour, 2-hour, 4 hour and 8 hour post calcein treatment using TIRF microscopy.

### AAV2 transduction efficiency assay

For transduction efficiency assays, cells were either left untreated or pretreated with 20 µM parthenolide for 2 hours prior to infection. Cells were then infected with AAV2 encoding a GFP transgene at an MOI of 5×10^3^ vgs/ cell or 2×10^4^ vgs/cell, in serum-free media for 3 hours. Following infection, the media was replaced with FBS-containing media, and cells were incubated for 48 hours before fixation and imaging. For flow cytometry analysis cells were untreated or pretreated with20 µM parthenolide for 2hrs before infection and then infected with AAV2 at 5×10^3^ MOI for 48 hours. GFP expression was quantified by flow cytometry (BD bioscience Accuri C6 plus) using 488 nm laser and 533/30 nm band pass filter.

### Immunofluorescence

Cells were fixed with either 4%PFA (15710, Electron Microscopy Science) or 3% PFA=+0.2% GA (Millipore) for 10 min at room temperature (RT) and washed with 1x PBS (Gibco). A NaBH_4_ solution (1 mg/ml) in PBS was used to quench the autofluorescence of GA. Cells were permeabilized and blocked with 3% BSA and 0.2% TritonX100 in PBS for 2 hours at RT. Cells were then incubated in primary antibodies for detyrosinated microtubule (ab48389; 1:100) or tyrosinated tubulin (ab6160; 1:100) for 2 hours at RT, or AAV2 (Fitzgerald 10R-A110a 1:100), or Phospho-GSK-3β (Ser9) (CST 5558; 1:200), or CLASP2 (Thermo Fisher PA5-109547; 1:100) overnight at 4° C, and then washed three times with washing buffer (0.2% BSA and 0.05% TritonX100) for 5min each. Then incubated with dye-tagged secondary antibody, donkey anti-rabbit (Jackson ImmunoResearch 711005152; 1:100), donkey anti-mouse (Jackson ImmunoResearch 715005150; 1:100), donkey anti-rat (Jackson ImmunoResearch 712005153; 1:100) in blocking buffer for 1.5 hr. followed by three times wash with PBS for 5min each. For STORM imaging secondary antibodies were labelled with Alexa Fluor-647 and Alexa Flour 405 at ∼1:4 ratio per antibody.

### Microscopy techniques

Confocal Images were acquired using a Zeiss laser scanning confocal microscope (LSM 780 system) equipped with 63X oil immersion, 40X oil immersion and 20X objectives. Argon laser 488 nm was used to excite GFP, DPSS laser 561 nm was used to excite Cy3/ AF 560 and HeNe laser 633 nm was used to excite the AF647 stained samples.

The 3D super-resolution Structured Illumination Microscopy (SIM) imaging was performed on fully motorized lattice SIM, ZEISS Elyra 7 (Carl Zeiss Ltd.) system equipped with Plan-Apochromat 63x/1.4 oil immersion objective lens, sCMOS camera and four filter sets with precisely mounted ACR-coded filter modules. 488 nm laser was used to excite the AF 488, 561 nm laser was used to excite the AF 560, and the 642 nm laser was used to excite AF 647 stained samples on a XY Piezo Scanning Stage. Image recoding was done on leap mode for the 3 times faster imaging, and the processing of the images was done by the in-built processing software of the Lattice SIM system.

TIRF microscopy was performed on a custom-built setup with a Nikon Ti2E microscope body, 1.49 NA 100x oil immersion objective lens, emission filter (ET705/72m; Chroma. Images were captured by an EM-CCD camera at an exposure time of 100ms per frame for 1000 frames for live-cell imaging and 10 frames for fixed cell imaging.

A few images were acquired using SAFeRedSTORM module (Abbelight) mounted on an Evident/Olympus IX3 microscope equipped with an oil-immersion 100x objective, 1.5NA (Evident/Olympus) and fiber-coupled 642 nm laser (450mW Errol). Images were collected with an ORCA-Fusion sCMOS camera (Hamamatsu). Image acquisition and microscope control were driven by 586 Abbelight’s NEO software.

### Correlative live-cell and STORM imaging

The live-cell imaging of AAV2-Cy3 was performed on the home-built TIRF microscope described above. Images were acquired for ∼1000 frames at 100 millisecond exposure followed by *in-situ* fixation of the cell. Cells were then immunostained for detyrosinated microtubule and the super resolution STORM image was acquired. 647 laser (MPB communications) was used to excite the reporter dye – Alexa fluor 647 (Invitrogen) and a 405 laser (Obis; Coherent) was used to excite the 405-activator dye (Invitrogen). The emission of the light was collected by Nikon 100x oil immersion TIRF-SR objective (NA 1.49) and laser quad band set with emission filter (TRF89902-EMET-405/488/561/647 nm laser Quad Band Set for TIRF application; Chroma). Image localizations were captured by EM-CCD camera at an exposure time of 20 milliseconds per frame for more than 60000 frames for each STORM image. STORM images were analyzed and rendered as previously described, using custom-written software (Insight3, provided by B. Huang, University of California, San Francisco, CA, USA ^64^).

To achieve channel registration, live-cell videos of AAV2 (Cy3 emission) excited with a 560-nm laser line and captured using a quad-band filter set were aligned with super-resolution images of detyrosinated microtubules (Alexa Fluor 647 emission) excited at 647 nm and acquired using the same filter set, following previously established methods ^27,28^. Briefly, to account for sample drift during imaging, a rigid shift along the x and y axes was determined using TetraSpeck beads (Thermo Fisher Scientific, USA) placed on the imaging dish glass. These beads were visible in both the live-cell AAV2 images and the STORM images of detyrosinated microtubule. The calculated shifts were then applied to the raw localizations from the super-resolution imaging, using custom algorithms as described previously ^27^.

### Single particle tracking

For single particle tracking AAV2 particle positions were tracked using custom-made, semi-automated particle tracking software. The x and y coordinates of the trajectories were extracted by fitting a 2D Gaussian function to the point spread function of the objects. To identify the active and passive transport phases, a MATLAB-based custom script was utilized, as described in earlier studies ^27^. Briefly, the trajectory analysis involved a moving window approach, using four-point segments along the coordinate data points. For each segment, the ratio of the displacement between the segment’s starting and ending points to the sum of displacements between each points in the segment was computed. Since segments overlapped, individual trajectory points contributed to multiple segments, and the ratio for each point was averaged across all its associated segments. This ratio served as a measure of linearity, with values near 1 indicating active transport. A threshold ratio of 0.7 was set to differentiate between active and passive phase. To refine the classification further, an angle-based criterion was applied: segments where consecutive displacement vectors formed angles smaller than 90° were labeled as passive. Additionally, mean square displacement (MSD) analysis was used to validate the classification, where active transport exhibited α value greater than 1.5. Only trajectories with at least five data points were considered for further analysis. Metrics such as run length, speed, pause frequency per trajectory, and pause duration were calculated from the processed data.

### *In vivo* combination therapy

For *in vivo* studies we utilized the prodrug DMAPT (Abcam; ab146189) which is a fumarate salt of parthenolide with improved bioavailability. The animals were divided into mock, DMAPT only, scAAV2 LP1-h.FIX, and Combination (DMAPT + scAAV2 LP1-h.FIX). Animals used in the study were 7–11 weeks old, with 5-7 animals per group. Animals in the DMAPT-only and Combination groups were pre-treated with the DMAPT (20µg/mouse) *via* oral gavage for 5 days before and one week after injection. The scAAV2 LP1-h.FIX and combination groups received a vector dose of 5 × 10¹⁰ vector genomes (vgs). Immediately before injections, the animals in the DMAPT-only and Combination groups received another dose of the drug. Post-injection, DMAPT was administered on alternate days for up to 7 days. Blood samples were collected after 30 days using retro-orbital bleeding into tubes containing 3.8% sodium citrate (as an anticoagulant). Plasma was isolated using standard methods and stored at –80°C until analysis. FIX activity was measured using the Hyphen chromogenic FIX assay (Hyphen BioMed, Neuville-Sur-Oise, France), following the protocol provided by the manufacturer. Calibrators, control reagents, and buffers were brought to room temperature for 30 minutes before resuspension. They were then resuspended using Milli-Q water, mixed gently, and left undisturbed for 30 minutes. Standards were prepared by serial dilution, ranging from a high concentration of 200% to a low concentration of 3.125%. Sample dilutions were performed using the buffer provided with the kit. Readings were taken at 450 nm, and a standard curve was generated by plotting a straight line of log (OD) versus log (Concentration).

### Immunohistochemistry

After 30 days of vector administration, animal liver tissue was harvested followed by blood collection. Liver tissue was fixed in 4% PFA in PBS overnight at 4° C. Sucrose gradient treatment was performed with 15% and 30% sucrose for 12 hours and 2 hours respectively. Tissue samples were further mounted in the PolyFreeze (Sigma Aldrich, Missouri, USA) and cryo-sectioning was done (Leica CM1520, Leica Biosystems, Wetzlar, Germany) to obtain the 10µm thick liver tissue sections. For immunohistochemistry, the tissue was then fixed again for 15 mins in 4% PFA. The blocking was done using 5% normal donkey serum and 0.2% tritonX100 in PBS for 2 hours at RT. Primary antibody staining was performed against detyrosinated microtubule (1:100), tyrosinated microtubule (1:100) and Factor-IX protein (1:250) overnight at 4° C. Secondary antibody (1:500) incubation was done for 2 hours at RT after PBS wash. This was followed by the nucleus staining with diamidino-2-phenylindole (DAPI) (Sigma Aldrich) and slides were then mounted with Prolong Gold (Thermo Fisher Scientific, USA).

### Image processing and quantification

SIM images were processed using FIJI ^65^. Maximum intensity projection was applied to all images, and brightness and contrast were adjusted uniformly across all conditions to ensure consistent representation and quantification of microtubule-PTM ratio data. For intensity quantification, image thresholding was performed using identical parameters for all conditions, and the integrated density was measured. The fluorescence-integrated densities of tyrosinated and detyrosinated microtubule were summed to determine the total microtubule density. The percentage of detyrosinated and tyrosinated microtubule was then calculated by dividing their respective integrated densities by the total microtubule density. To quantify the percentage of AAV2 associated with detyrosinated microtubule, multiple frames from each z-stack were analyzed. The percentage was calculated by dividing the number of AAV2 particles colocalized with detyrosinated microtubule by the total number of AAV2 particles. For track displacement analysis we utilized the TrackMate ^66,67^ plugin from FIJI. For zonal distribution analysis, the outer boundary and nucleus of the cells were first outlined. Straight lines were then drawn from multiple points along the nuclear boundary to the outer boundary. Each line was divided into three equal segments, and corresponding boundaries were drawn to define three distinct regions: the outer zone (adjacent to the plasma membrane), the mid-zone (central cytoplasm), and the inner zone (proximal to the nucleus). The number of AAV2 particles within each zone was manually quantified. For Rab7-AAV2 colocalization analysis, the colocalization of Rab7 with AAV2 was manually scored. For endosomal escape analysis using the calcein assay, the integrated density was measured within the cytoplasmic area devoid of puncta, and values were compared across different conditions after subtracting the background integrated density.

## Statistical analysis

All the data was analyzed for significance using a two-sample *t*-test and two-way ANOVA for *in vivo* studies. When required data was normalized to a maximum value. The *p*-value of ≤ 0.05 was considered statistically significant. All the analysis was performed using OriginPro 2021b (Academic).

## Data availability statement

The data set studied and investigated during the current study are accessible from the corresponding authors (N.M. and G.R.J) upon reasonable request.

## Acknowledgments

We thank Prof. Melike Lakadamyali (University of Pennsylvania, Philadelphia, PA) for supporting the research with reagents, software. We thank Indranil Banerjee (IISER Mohali, India), Carsten Janke (Institut Curie, France) and Minhaj Sirajuddin (InStem, Bengaluru, India) for providing plasmids. We thank Prof. Appu Kumar Singh (IIT Kanpur) for critically reading the manuscript. NM thanks Prof. Pradip Sinha (IIT Kanpur) for suggestions and mentorship.

S.T. expresses gratitude towards the Council of Scientific and Industrial Research, India for providing financial assistance during research. N.M. and G.R.J. acknowledge funding from the Indian Council of Medical Research, India (ICMR F.No. 2021-15239/GTGE/Adhoc-BMS) and the Indian Institute of Technology, Kanpur, India. The authors gratefully acknowledge the contribution of Jeganath A., Deepak M. Khushalani, and Sanket Patil in helping establish the laboratory facility at IIT Kanpur, India. The authors also acknowledge Jemina J for the help with single particle tracking. Furthermore, Pratiksha Sarangi has been noted for her help during animal studies. Images were partially created using <u>BioRender.com</u>.

## Author contribution

S.T. designed and carried out the experiments and performed the data analysis. S.H. contributed to *in vivo* experiments. J.K. contributed to in vitro experiments. D.C did the experimental analysis of Hemophilia samples. N.M. and G.R.J. designed and supervised the research. S.T., N.M., and G.R.J. wrote the manuscript. All authors provided feedback on manuscript.

