## Supplementary Figures for "Suppressing microtubule detyrosination augments AAV2 endosomal escape and gene delivery"

#### Figure S1

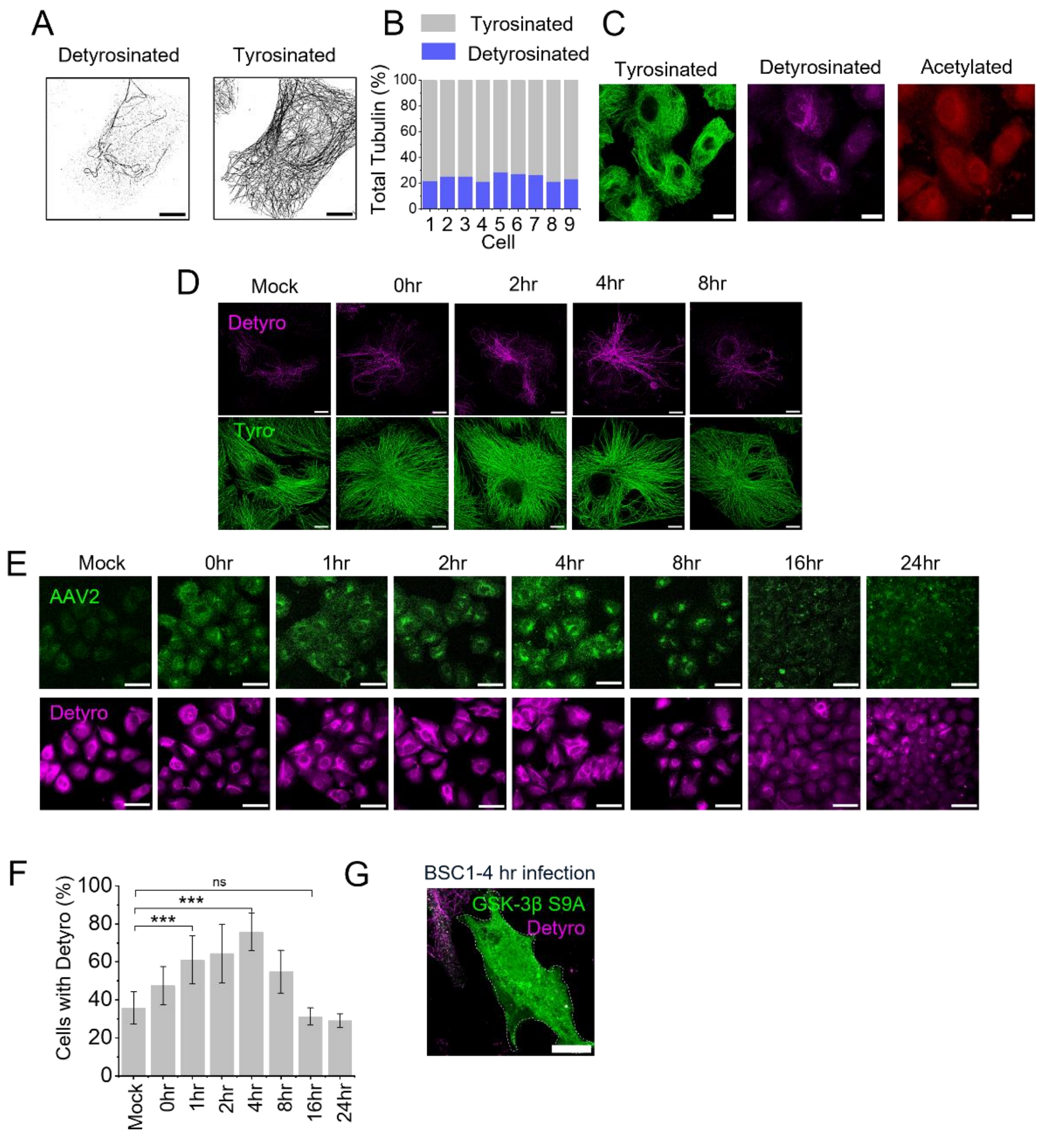


**Supplementary figure 1. Microtubule PTM quantification during AAV2 infection.**
**(A)** SIM images with intensity threshold-applied to enumerate the integrated density of tyrosinated and detyrosinated microtubules (see Methods) and quantify the percentage of microtubule-PTM in each Huh7 cells. **(B)** Bar plots depicting the proportion of tyrosinated (grey) and detyrosinated (blue) tubulin in individual cells, at 4-hour infection **(C)** Confocal microscopy of Huh7 cells infected with AAV2 for 4 hours and immunostained for tyrosinated (green), detyrosinated (magenta), and acetylated (red). Note that Huh7 cells do not show detectable levels of acetylated microtubules as reported previously [9,10]. **(D)** Structured illumination microscopy (SIM) images of BS-C-1 cells infected with AAV2 and fixed at 0-, 2-, 4-, and 8-hours post-infection show the level of detyrosinated (magenta) and tyrosinated (green) microtubules. **(E-F)** Confocal imaging of mock and AAV2-infected Huh7 cells fixed at 0-, 1-, 2-, 4-, 8-, 16-, and 24-hours post-infection reveals an increase in the number of cells with microtubule detyrosination over time. Quantification of the percentage of Huh7 cells expressing detyrosinated microtubule at different time points post-infection (F) (n=12 cells for each time point) highlights temporal modulation of tubulin post-translational modifications**.** **(G)** Suppression of detyrosinated microtubules observed in BSC-1 cells expressing GSK3β S9A, indicating that the signaling effect is not cell-type specific. **(H)** Overexpression of wild-type CLASP2 in Huh7 cells followed by mock or AAV2 infection. CLASP2 is associated with microtubules only in infected cells, indicating infection-specific activation. Bars represent the mean value and whiskers represent standard deviation. Statistical analysis via two paired two-sample t-tests confirmed significant differences (***p<0.001). Scale bars: (A, F, G and H) 10 µm, (C) 20 µm and (D) 50 µm.

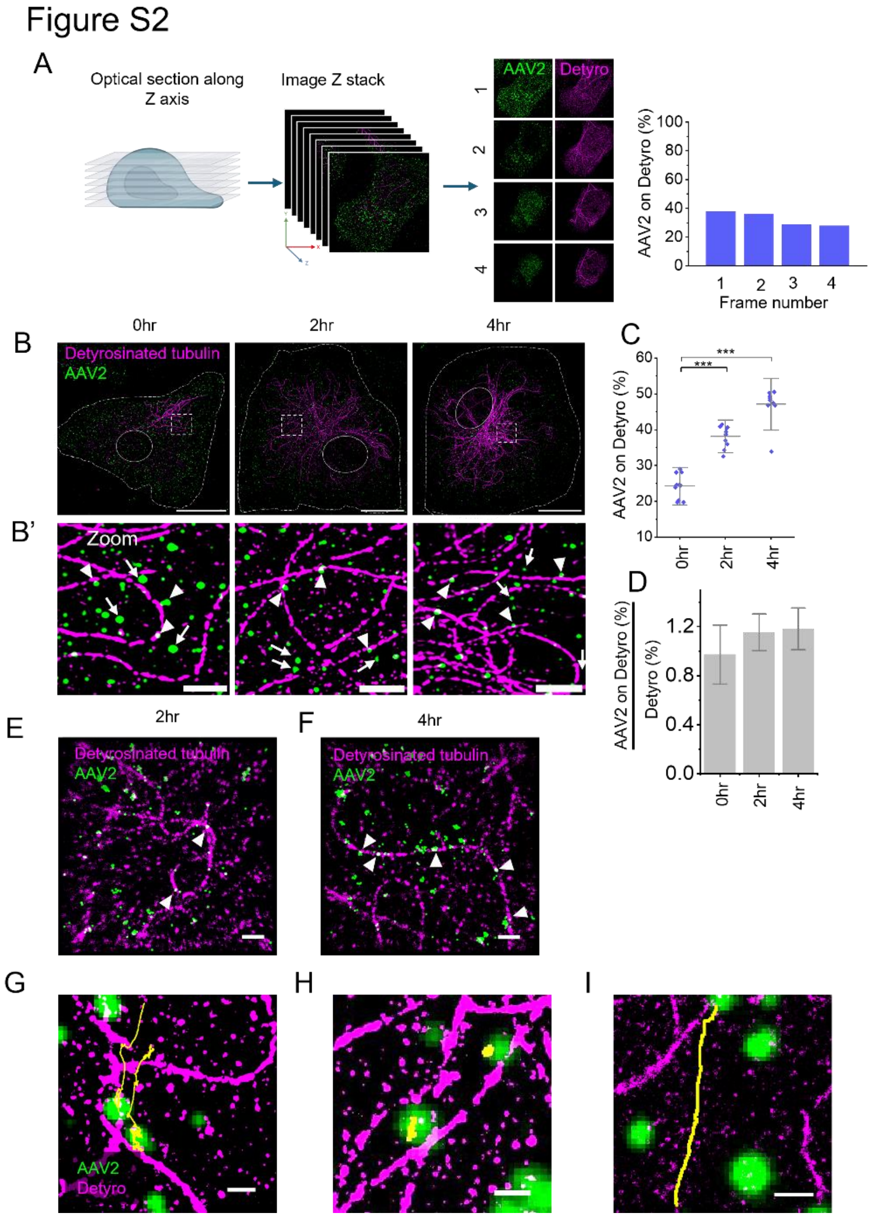


**Supplementary Figure 2. Quantitation of AAV2 colocalization with detyrosinated tubulin.**

**(A)** Schematic representation of optical sectioning and z-stack image acquisition for quantifying AAV2 localization on detyrosinated tubulin, from a SIM z-stack image of AAV2 (green) and detyrosinated tubulin (magenta). From each Z-section the percentage of AAV2 localized on detyrosinated microtubule was enumerated. **(B-D)** SIM images of BS-C-1 cells infected with AAV2 and fixed at 0-, 2-, and 4-hours post-infection indicating zoomed-in regions (B') from (B) highlight specific instances of AAV2 localization with arrowhead and arrows show AAV2 localized and not localized to detyrosinated microtubules, respectively. The graph shows the colocalization (C) and enrichment (D) of AAV2 on detyrosinated microtubules over time. **(E-F)** STORM imaging of Huh7 cells infected with AAV2 shows the association of AAV2 (green) with detyrosinated tubulin (magenta) at 2 hours (E) and 4 hours (F) post-infection, with arrows indicating AAV2 particles on detyrosinated tubulin. **(G-I)** Images demonstrate the last frames of the SI-videos (Video 1-5) with overlay of AAV2 (green) tracks and STORM imaging of detyrosinated tubulin (magenta) videos. The dynamic behavior of AAV2 is captured in real-time (1-hour post-infection), with its trajectory overlaid on the video as a track (yellow), illustrating the path of viral particles as they interact with the microtubule. Examples from the the last frame of video show that viral motility is hindered when encountering detyrosinated tubulin (G). The virus particles remain hindered on detyrosinated tubulin until the end of the recording (H). Directed transport of AAV2 along microtubule regions other than detyrosinated tubulin remains unhindered, allowing faster movement (I). Scale bar: (B) 10 μm, (B') 2 μm and (E-I) 1 μm.

### Figure S3

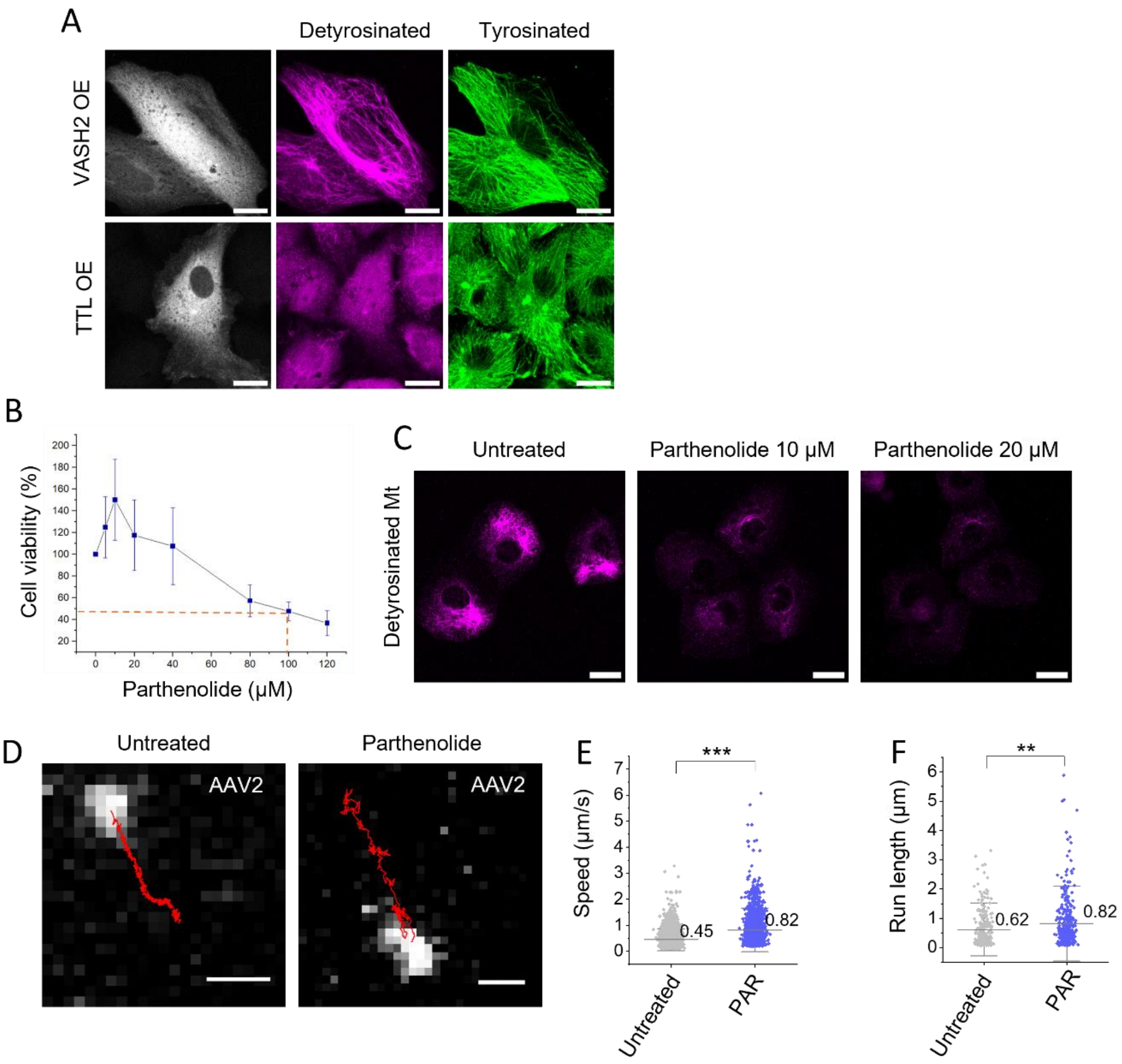


**Supplementary Figure 3. Modulating detyrosination/tyrosination levels and their impact on AAV2 trafficking.**

**(A)** Confocal images of Huh7 cells overexpressing VASH2 or TTL with corresponding detyrosinated and tyrosinated tubulin immunostaining. VASH overexpression shows a significantly high level of detyrosinated tubulin in Huh7 cells and TTL overexpression suppresses the detyrosination and upregulates microtubule tyrosination. **(B)** Concentration dependent cytotoxicity of parthenolide was assessed using an MTT assay- and $IC_{50}$ was analysed ($IC_{50}$= 100 μM). The graph shows mean values and standard deviation (n=3). **(C)** Imaging untreated and parthenolide-treated Huh7 cells (10 μM and 20 μM) infected with AAV2 reveals drug-induced suppression of detyrosinated microtubules. **(D)** Single-particle tracking of AAV2 in parthenolide-treated cells shows impaired viral motility compared to untreated control. **(E-F)** Graphs quantify AAV2 speed (E) and run length (F) under untreated and parthenolide-treated conditions, highlighting the effects of microtubule modification on viral dynamics. Statistical analysis showing **$p<0.005$ and ***$p<0.001$. Scale bars: 20 μm (A, C) and 1 μm (D). $IC_{50}$, half-maximal inhibitory concentration; MTT, 3-(4,5-dimethylthiazol-2-yl)-2,5-diphenyltetrazolium bromide.

Figure S4

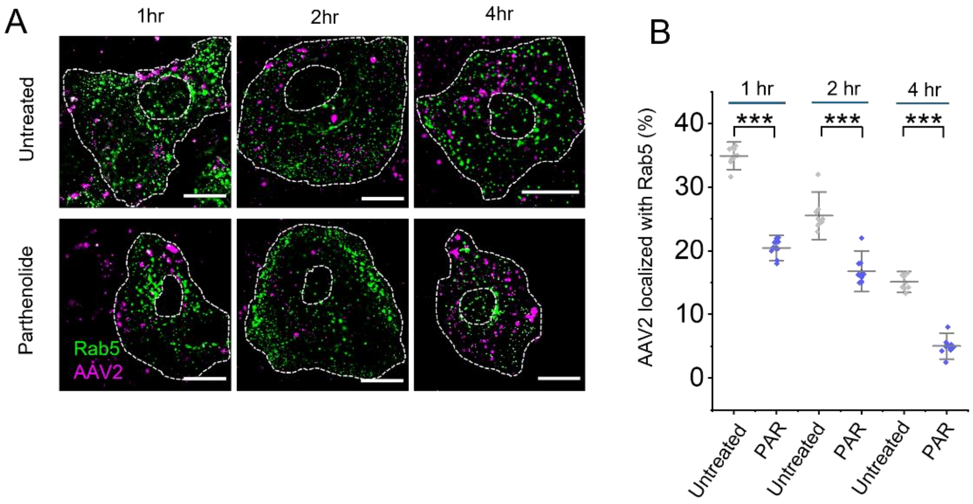


**Supplementary Figure 4. Endosomal sorting of AAV2.**

**(A-B)** Confocal imaging and Quantification of AAV2 (magenta) colocalization with early endosomes decorated by Rab5-GFP (green). Parthenolide-treated cells show significantly less colocalization of AAV2 with Rab5 compared to untreated controls (n=10) (B). In the graphs, the grey line indicates mean, and whiskers show the standard deviation from ten cells analyzed (n=10). Statistical significance assessed via a two paired two-sample t-test. Significance indicated as ***p<0.001. Scale bars: (A) 20 μm.

### Figure S5

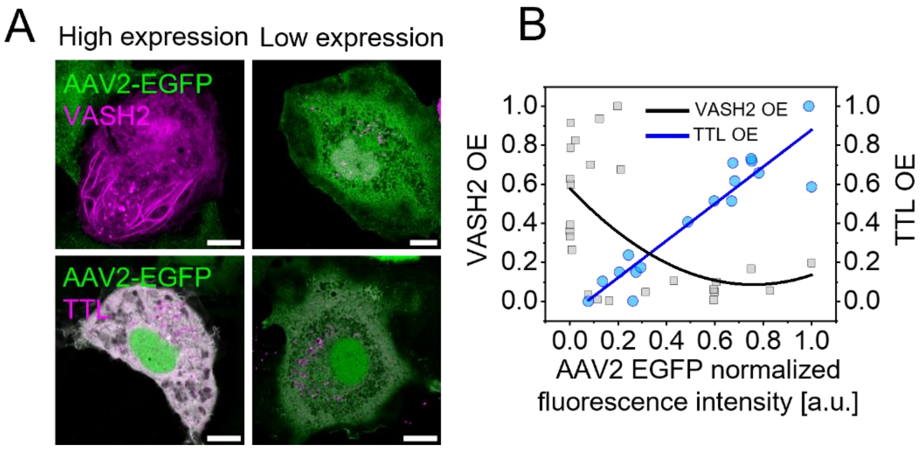


**Supplementary Figure 5. VASH/ TTL overexpression (OE) and correlation with AAV2 transgene expression.**

**(A-B)** Confocal imaging shows the effect of differential levels of VASH or TTL expression on AAV2 -EGFP gene transduction (A) and their quantification (B). Scale bars: (A) 10 µm.
